# Aerobic carbon metabolism modulates nitrite ammonifiers for inhibiting nitrogen loss as revealed by microcosm experiment of agricultural upland soil

**DOI:** 10.1101/2024.11.04.621907

**Authors:** Xiaogang Wu, Siyu Yu, Weikang Sui, Xinyu Zhang, Ji Li, Qiaoyu Wu, Xiaojun Zhang

## Abstract

Denitrification and dissimilatory nitrate reduction to ammonium (DNRA), two divergent nitrogen metabolism pathways, share nitrite as a common intermediate during anaerobic reduction. However, these pathways influence the environment and soil nutrients differently. Unlike denitrification, which involves nitrogen loss through gaseous nitrogen emission and nitrate leaching, DNRA is beneficial for the conservation of nitrogen. Nevertheless, the key mechanism underlying the joint regulation of aerobic carbon metabolism to nitrite ammonifiers in soil is still unclear, although previous studies have reported factors such as carbon and oxygen can independently affect DNRA. Here, microcosm experiments with agricultural upland soil were conducted under different aeration conditions supplemented with labile carbon to analyze the process of anaerobic nitrogen metabolism. The results indicated that denitrification exclusively dominated nitrite reduction when the soil was directly placed in an anaerobic environment. Nonetheless, a significant increase in DNRA activity and the attenuation of denitrification were detected when the soil was incubated aerobically with the addition of glucose prior to anaerobic incubation. Specifically, up to 55.8% of nitrite reduction switched to nitrogen conservation mainly via DNRA under high-carbon conditions. Quantitative assays of the *nrfA* gene and metagenomic data revealed a significant increase in DNRA-related genes after aerobic carbon metabolism. Furthermore, *nrfA* gene sequence analysis revealed a significant shift in the composition of nitrite ammonifiers community. These results indicate that nitrate/nitrite metabolic flux in the soil could be regulated to enhance DNRA by stimulating facultative anaerobic nitrite ammonifiers such as *Sedimentibacter* under alternating aerobic and anaerobic environments with carbon metabolism.

**IMPORTANCE:** This study revealed that aerobic conditions with a carbon supply strongly influence the community assembly of nitrite ammonifiers. Nitrite ammonifiers such as *Sedimentibacter* enriched under sufficient carbon and aeration conditions promoted the subsequent DNRA process under anaerobic conditions. Therefore, regulating the competition between DNRA and denitrification by altering functional microbiota would be a promising approach for improving the efficiency of fertilizer application and for reducing greenhouse gas emissions from denitrification. These findings are essential for understanding the biological mechanism of nitrogen cycling in dryland agricultural soil. Based on these findings, we may design a strategy to increase the abundance of nitrite ammonifiers in agricultural soil, which might promote DNRA metabolism to improve nitrogen retention. This strategy would ultimately enhance fertilizer use efficiency and mitigate the detrimental environmental effects of nitrogen fertilizer abuse.

## INTRODUCTION

Nitrate and nitrite are important forms of inorganic nitrogen in agricultural soil, especially in fertilized upland fields, where nitrification is active due to sufficient ammonium and good aeration. Most soil encompasses very diverse niches with different nutrients and physiochemical properties, such as both aerobic and anaerobic environments. Under anaerobic conditions, the primary metabolic pathway for nitrate is microbial reduction, with nitrite as an intermediate product, which is a hub intermediate in the reductive metabolism of inorganic nitrogen. In the soil environment, different microbial groups compete for available nitrate/nitrite through denitrification, anaerobic ammonium oxidation (anammox), and dissimilatory nitrate reduction to ammonium (DNRA) (1). Unlike the other processes that convert inorganic N species to N_2_, DNRA reduces nitrite to ammonium and is beneficial for the conservation of bioavailable nitrogen and enhancing fertilizer use efficiency (2, 3). Thus, the conversion of nitrate to ammonium is conducive to the retention of nitrogen in agricultural soil systems. These branched nitrogen metabolic pathways are individually selected as the main or partial stream for further reduction depending on the biotic or abiotic conditions in situ (4, 5). Consequently, these conditions determine the quantity and form of nitrogen in agricultural systems. Furthermore, abiotic or biotic conditions determine the availability of readily usable nitrogen for assimilation into crops (6). Therefore, studying the mechanism by which nitrate- and nitrite-reducing metabolism are regulated is important for both agricultural economy and environmental protection. Promoting the occurrence of DNRA in overfertilized arable soil, especially in upland farm soil, would be beneficial for reducing soil nitrogen loss and mitigating contamination of the aquatic environment by nitrate leaching. Switching the metabolism of nitrate/nitrite reduction to DNRA rather than denitrification will also greatly mitigate nitrous oxide emissions as previously described (7).

DNRA has been widely reported in soils of different terrestrial ecosystems, such as wetlands, forests, grasslands, paddy soils, and other farmlands (8). DNRA could be an important process in some circumstances; for example, in temperate rainforest soil in Chile, almost all nitrate is consumed by DNRA (9, 10). However, DNRA has been shown to account for only 5-19% of nitrate reduction in temperate freshwater environments, such as riparian fens and wetlands (11–15), and has a similar range in paddy soils (16). Denitrification is recognized as the main bacterially facilitated process of nitrate and nitrite reduction in most circumstances (1, 17, 18). DNRA is often considered negligible in croplands (19, 20). Many studies have reported the relationship between DNRA and denitrification. For example, DNRA and denitrification compete with each other because both nitrate and nitrite are common substrates of these two pathways (4, 21). The addition of labile carbon to the soil usually leads to a strong “priming effect”, consequently promoting the metabolism of nitrogen due to the increase in electron donors, such as the reduction of nitrate under anaerobic conditions (22, 23). Accordingly, the supply of carbon is important for both DNRA and denitrification (22, 23). However, studies have shown that DNRA becomes more competitive during the reducing state with a high dose of fermentable dissolved organic carbon (DOC) (5, 16, 24–28). DNRA microorganisms are enriched under nitrate-limiting conditions and are outcompeted by denitrifiers under fermentable DOC limitation (4, 21). DNRA activity and the DNRA population were significantly positively correlated with the carbon-to-nitrogen (C/N) ratio (4, 29). Therefore, the C/N ratio might be a key factor in regulating soil DNRA and denitrification (29). This was also proven in experiments with pure bacterial cultures. In a study, an isolate of *Shewanella* that possessed both denitrification and DNRA metabolic functions showed that its metabolic flux on DNRA and denitrification could be regulated by the C/N ratio (30). However, a high C/N ratio does not always guarantee a high ratio of DNRA for denitrification. For example, the denitrification capacity was highest in a grazing grassland area of a floodplain where the lability of organic carbon was significantly greater than that in other ungrazed areas, and the denitrification capacity was an order of magnitude greater than that of DNRA (20). The interactions of multiple factors underlie the competition between these two pathways, for which the C/N ratio may not be the most critical factor.

Previous studies have repeatedly reported that controlling the C/N ratio could regulate the activity of denitrification and DNRA (5, 16, 24–28). However, the results of microcosm experiments using soil from the North China Plain are not consistent with the perspective that a higher C/N ratio enhances DNRA activity. Denitrification still dominates nitrate and nitrite reduction under directly anaerobic conditions with the addition of a high dosage of fermentable DOC. According to the analysis, the increased activity of the denitrifying microbiota controlled the flow of nitrogen metabolism pathways (31). Therefore, it is speculated that dryland soil does not possess greater DNRA activity under directly anaerobic conditions. Previous studies have indicated that increased DNRA activity is associated with increased levels of functional DNRA genes, such as *nrfA,* in sediments (32). The DNRA potential rates were affected by altering the community structure of the DNRA bacterial guild (33, 34). Hence, the DNRA functional microbiota together with the C/N ratio determine the DNRA rate and its ratio to denitrification in the reduction of nitrate. The enrichment of the DNRA functional microbiota provides a foundation for improving DNRA activity. However, under directly anaerobic conditions, DNRA activity did not increase with a high dosage of fermentable DOC in the soil from the North China Plain (31). The enrichment strategy still needs further investigation (35).

According to previous findings, oxygen is also an important regulator between denitrification and DNRA (36, 37). Using sediments in the southern basin of Lake Lugano, relatively high O_2_ tolerances for DNRA and denitrification were observed (70 µmol L^−1^), and DNRA was systematically favored over denitrification at relatively high O_2_ levels (36). According to research on sediments on a global scale over six continents, riparian zones, which are typical environments of oxic-anoxic interfaces, have a high rate and abundance of DNRA (38). These results suggest a correlation between DNRA function and aeration conditions. Biochemically, nitrate-reducing bacteria can reduce nitrate to nitrite, which is catalyzed by two different nitrate reductases, membrane-bound nitrate reductase (NAR) and periplasmic nitrate reductase (NAP) (39, 40). Nitrite is the nodal intermediate metabolite for both metabolic pathways in denitrification and DNRA, and it can be directly reduced to ammonium by nitrite reductase (encoded by *nrfA*) of nitrite ammonifiers in the DNRA pathway (41, 42). However, denitrifying bacteria perform denitrification anaerobically by reducing nitrite to gaseous nitrogen via the use of NIR, NOR, and NOS (encoded by *nirS*/*nirK, norB and nosZ*). It has also been proposed that NAP facilitates the conversion of aerobic respiration to anaerobic metabolism (43). Studies on enzyme expression in *Paracoccus pantotrophus* and *Paracoccus denitrificans* have revealed that NAR is predominantly expressed under anaerobic denitrifying conditions, while NAP is predominantly expressed under aerobic growth conditions (44). Compared with NAR-containing bacteria, bacteria containing NAP might have advantages in an environment that often experiences a transition of oxygen conditions. A higher C/N ratio cannot enhance DNRA activity in North China Plain soil (31). Hence, oxygen and DOC might jointly influence ammonifiers in the DNRA process on the North China Plain via aerobic metabolism. However, little attention has been given to the effects and mechanism of aerobic carbon metabolism on the ecology of nitrite reduction.

In this study, a soil microcosm experiment was performed to investigate the effects of aerobic carbon metabolism on soil nitrite ammonifiers and to attempt to develop an enrichment strategy for the nitrite ammonifiers. The soil was collected from arable land on the North China Plain (NCP, Table S1), where a dry climate and long-term intensive nitrogen fertilization characterize agricultural farmland (45–48). Since NCP soil possessed the relatively low concentration of original ammonium (Table S1), we only compared DNRA and denitrification without anammox in this study. We analyzed the nitrite ammonifiers in typical NCP soils under different conditions of aeration and fermentable DOC addition (Fig. S1). Quantitative PCR was performed to evaluate the abundance of nitrite ammonifiers, while amplicon and metagenomic sequencing were performed to analyze the composition of the functional microbiota. The present research aimed to develop a practical strategy for increasing the ratio of DNRA to denitrification during nitrate reduction and to elucidate the biological mechanism regulating nitrate-reducing metabolism, where nitrate/nitrite often appears to be the main component of inorganic nitrogen.

## MATERIALS AND METHODS

### Soil samples

Typical NCP agricultural upland soil was collected from cropland at Quzhou Experimental Station (36° 86’N, 115° 02’E), Hebei Province, northern China. The soil in the farm field is calcareous fluvo-aquic soil (calcareous Cambisols according to the FAO Classification) with normal fertilization management (49). The soil cultivation type belonged to winter wheat-summer maize double-cropping. The soil samples were collected from the top 20 cm using a sterile hand corer (10 cm diameter). The soil was collected via multipoint sampling in the field and then placed in an ice box to be stored at low temperature and transported to the laboratory. Composite soil samples from numerous auger borings were sieved to 2 mm to remove stones, roots and crop residues and then stored at 4°C until the experiment began. The basic physiochemical properties of the investigated soil are shown in Table S1.

Before the designed incubation experiments, a pretreatment step was applied to the soil to activate the soil microorganisms and minimize the background nitrate/nitrite as much as possible from the soil. Briefly, the pretreatment mainly included the following steps. The soil samples were firstly incubated at 25°C for 7 days. Then, 30 g of fresh soil sample was added to a 100 ml serum vial, and the gas in the sealed vial was replaced with ultrahigh-purity helium using a pump ventilation system (Shanghai Jincheng Co., China). Then the samples were anaerobically incubated for 15 days at 25°C to reduce the nitrogen content. After preincubation, the soil acted as the baseline sample for subsequent experiments, including microcosm incubation and isotopic tracer experiments. Physiochemical parameter determination and microbiome analysis were performed as described for the untreated control (UNC). The inorganic nitrogen and DOC contents of the soil samples were determined as previous study (31). Here, the determination methods were also briefly described as followed. NO_3_^-^-N was extracted using 1 M KCl solution and then analyzed using an automatic chemical discontinuity analyser. NO_2_^-^-N was determined by N-(1-naphthyl)-ethylenediamine dihydrochloride spectrophotometric method. NH_4_^+^-N was estimated by the indophenol blue method. DOC was monitored by a total organic carbon analyser (TOC-LCSH, Shimadzu, Japan).

### Experimental setup

Since nitrite is the direct substrate common for both metabolic pathways in denitrification and DNRA, using nitrite as the substrate of the experimental microcosm could exclude other interfering factors when exploring the two metabolic flows (Fig. S1). The total incubation time of aerobic and anaerobic period for NCP agricultural upland soil was set according to previous studies (49–51). Glucose was added as carbon source for the microcosm experiment according to previous studies (31, 50). Aerobic soil incubation experiments were conducted to modulate the soil microbiome at 25°C for 7 days in serum vials, each containing 30 g of UNC baseline soil amended with 1000 mg·kg^-1^ soil glucose. The headspace gas in the sealed vials was replaced with mixed gas (21% oxygen and 79% helium) every day using a pump ventilation system, which was named as AEI treatment. Nine replicate vials were used in the AEI treatment. Three of them were subjected to destructive sampling and used for the analysis of physiochemical parameters and the microbiome of the AEI treatment group. The other six vials were used for subsequent treatments. The average DOC content in these vials in the AEI treatment group decreased to nearly 200 mg·kg^-1^ soil after incubation. First, after the AEI treatment, all six vials were supplemented with sodium nitrite (60 mg NO_2_^-^-N kg^-1^ soil). Then, three of them were injected with glucose (final concentration equivalent to 1000 mg DOC·kg^-1^ soil), which was named as AEH (pre-aerobically incubated soil with high carbon) treatment. Another three vials were injected with sterilized water as control, which was named as AEL (pre-aerobically incubated soil with low carbon) treatment. Both the AEL and AEH treatments were incubated in the ROBOT system (robotized incubation system for monitoring gases) (52) for 7 days under anaerobic conditions maintained with helium in the headspace at 25°C in triplicate. The dynamics of gaseous nitrogen production were monitored by the ROBOT system.

The microcosm experiments involving direct anaerobic cultivation were also set up in 100 ml serum vials, each containing 30 g of fresh UNC soil that was amended with sodium nitrite as the nitrogen source (60 mg NO_2_^-^-N kg^-1^ soil, approximately 128.6 μmol per vial) and glucose as the carbon source at a concentration of 200 mg DOC·kg^-1^ soil (with a content equivalent to that of the AEL treatment) or 1000 mg DOC·kg^-1^ soil. The treatment with glucose in concentration of 200 mg DOC·kg^-1^ soil was named as ANL (anaerobic incubated soil with low carbon) treatment, and the treatment with glucose in concentration of 1000 mg DOC·kg^-1^ soil was named as ANH (anaerobic incubated soil with high carbon) treatment. The sample vials of the ANL and ANH treatments were incubated for 7 days under anaerobic conditions maintained with helium in the headspace at 25°C. All treatments were conducted in triplicate. The dynamics of gaseous nitrogen production were continually monitored by the ROBOT system.

### Calculation of soil nitrogen loss and conservation

The proportion of nitrogen loss via denitrification was calculated by the ratio of gaseous nitrogen products to the total contents of initial NO_3_^-^-N and NO_2_^-^-N. The remaining nitrogen, other than gaseous nitrogen and residual nitrate/nitrite, was attributed to nitrogen conservation in the soil microcosm. To analyze the complete DNRA process from nitrate to ammonium under anaerobic conditions for precluding the other ammonification, potassium nitrate (partially replaced with isotopic potassium nitrate (NO_3_^-^-^15^N)) was used as the substrate for reduction in further isotopic validation experiment. The method details of the validation experiment were similar to those of the AEL and AEH treatments, except for differences in substrate and incubation time. Specifically, the AEI soil was set up with both unlabeled (^14^N) and labeled nitrate either with or without glucose. The atomic ratio of ^15^N in labeled treatments is 9.24%. All of the treatments were incubated under anaerobic conditions at 25°C for 3 days in triplicate. The concentration of NH_4_^+^-^15^N kg^-1^ was determined after incubation. Based on methods described in the literature (53, 54), the NH_4_^+^-N in the soil samples was transformed into N_2_O, and the δ ^15^N content and the ^15^N/^14^N atomic percentage (atom%) were determined by a stable isotope ratio mass spectrometer (Thermo Fisher, USA) as previously described (19).

### Sequencing and quantification analysis of the *nrfA* gene in soil samples

The DNA of the soil samples was extracted from 0.5 g of soil, as described previously (55, 56). Then the extracted DNA from the soil samples was also used as a template to amplify the expected 250 bp amplicon of the *nrfA* gene. Touch-down PCR was performed as previously described (57). The reaction was adjusted according to two-step PCR for the construction of a sequencing library according to the protocol of the MiSeq system (Illumina, USA) provided by Illumina for amplicon sequencing (58). The thermal conditions of the first PCR step (amplicon PCR) were as follows: 95°C for 5 min, 95°C for 15 s, 58°C for 1/cycle, 52°C for 30 s, 72°C for 30 s for 7 cycles, 95°C for 15 s, 52°C for 30 s, 72°C for 35 s, 78°C for 15 s for 30 cycles, and 72°C for 10 min. The primers for amplicon PCR of the *nrfA* gene were nrfAF2aw+adapter (5’-TCGTCGGCAGCGTCAGATGTGTATAAGAGACAGCARTGYCAYGTBGARTA -3’) and nrfAR1+adapter (5’-GTCTCGTGGGCTCGGAGATGTGTATAAGAGACAGTWNG-GCATRTGRCARTC -3’). The thermal conditions of the PCR (index PCR) were as follows: 95°C for 5 min ×1, 95°C for 15 s, 52°C for 30 s, 72°C for 35 s, 78°C for 15 s ×7, and 72°C for 10 min. Sequencing of the PCR amplicons and quality control of the raw data were also performed as described previously according to the protocol for the MiSeq system.

Bioinformatics and sequencing data of the *nrfA* gene were obtained via QIIME2 (59). Sequence primers and adapters were removed, and the data were denoised by DADA2 (60). The denoising process included clustering sequences with 100% similarity, identifying chimeras and removing low-quality sequences. The amplicon sequence variables (ASVs) were obtained after denoising. This approach could increase the reliability and reproducibility of analyzing the amplicon sequences of functional genes (61). Analyses of alpha and beta diversity were performed with QIIME2 (59). Linear discriminant analysis (LDA) effect size analysis (LEfSe analysis) was performed to identify the ASVs that increased in response to high-dose DOC. The ASVs of AEI, AEL and AEH were compared with the ASVs of the UNC sample, and the ASVs were increased in all comparisons. The logarithm threshold of the LDA score was set as 2.0, and the analyses were performed with the LEfSe module on the website http://huttenhower.sph.harvard.edu/galaxy. These ASVs, both increased and dominant (relative abundance>1%), were recognized as the key ASVs. The taxonomy of the key ASVs was analyzed by BLASTX in the GenBank database as previously described (34).

Quantitative PCR was carried out on a Light Cycler 96 (Roche, Switzerland) using SYBR Green as a fluorescent dye to determine the relative abundance of the *nrfA* gene and 16S rRNA gene as previously described (62). The primers for the quantitative PCR of the *nrfA* gene were nrfAF2aw (5’-CARTGYCAYGTBGARTA-3’) and nrfAR1 (5’-TWNGGCATRTGRCARTC-3’) (62). The primers for the quantitative PCR of the 16S rRNA gene were Uni331F (5’- TCCTACGGGAGGCAGCAGT -3’) and Uni797R (5’- GGACTACCAGGGTATCTAATCCTGTT -3’) (63).

### Shotgun metagenomic sequencing and data analysis

Soil samples were collected for DNA extraction with the Omega Bio-Tek soil DNA kit and library construction using the Illumina TruSeq DNA sample preparation guide. To study the genes and functional bacteria in the samples subjected to different treatments in detail, the metagenomic DNA of twelve soil samples from the AEI, AEH, ANH and UNC groups, each in triplicate, was sequenced (2 × 150 bp) on an Illumina NovaSeq sequencer (by Personalbio, Shanghai). The data were quality-filtered by using fastp v0.20.0 software (64). To analyze the full-length protein sequences of the functional genes, contigs were generated by assembling high-quality reads using MEGAHIT (https://hkubal.github.io/megabox/) (65) with the default parameters --k_list 33, 55, 77, 99, 127. Using the software MMseqs2, the contig sequence set was deredundant with the linclust mode, according to the ratio of 95% similarity and 90% of the alignment region coverage of the short sequence, to obtain a nonredundant contig set. Open reading frames (ORFs) were predicted from all contigs using MetaGeneMark (http://exon.gatech.edu/GeneMark/) (66). The predicted ORFs were clustered to generate a nonredundant protein catalog (5,199,375 protein sequences in total). The nonredundant protein sequences were annotated by alignment with the NCBI-nr protein database by MMseqs2.

Approximately 0.01% of the genes (57562 in total) were assigned to N-cycle gene families according to BLAST analysis of the NCycDB database (67). The *narG* (nitrite reductase), *nirK*/*nirS* (nitrite reductase), *nosZ* (nitrous oxide), and *nrfA* (nitrite reductase-ammonia formation) genes were included as representative genes in the comparative analysis of denitrification and DNRA pathway-related genes across different treatments.

Pairwise comparison of soil samples under different culture conditions was performed to detect functional genes that were significantly more abundant in certain samples by setting thresholds of |log_2_FC| > 0, p value < 0.05 and fdr<0.05 in the program edgeR (68). Bacterial diversity based on metagenomic data was calculated according to the Shannon index, and difference analysis was performed using the Kruskal‒Wallis test with R software. The composition and abundance of the nitrite ammonifiers were determined via taxonomic annotation of the *nrfA* gene at the species level via metagenomic reads. Principal component analysis (PCA) was subsequently performed based on the nitrite ammonifiers abundance data. Permutational multivariate analysis of variance (PERMANOVA) was performed to compare the differences in the community structure of nitrite ammonifiers among the different treatments.

For phylogenetic reconstruction, sequences of the *nrfA* genes were aligned using MAFFT v7.505 (69), and the alignment was trimmed using trimal v1.4 (70). Subsequently, IQ-TREE(71) was used to find the optimal model for building a maximum-likelihood tree with 1000 bootstrap replicates. The Interactive Tree of Life (iTOL) online tool (https://itol.embl.de/) was used to visualize and annotate the phylogenetic tree and create the final figure. The LEfSe analysis method was used to identify differentially abundant microorganisms that can be used as potential biomarkers associated with DNRA. The criterion for the LDA score for discriminative features was >2.0, and the alpha value for the factorial Kruskal–Willis test among the classes was <0.05. All analyses carried out through LEfSe were performed through the Galaxy server. R version 4.1.3 was used for performing the statistical analyses and visualizing the results.

## RESULTS

### Soil nitrogen species and dissolved organic carbon after preincubation

After preincubation, the soil nitrogen species and DOC content were measured again as a baseline for subsequent experiments (Table S2). Background nitrate/nitrite has not been totally removed. The UNC soil NO_3_^-^-N content was 5.6±0.3 mg/kg, which was equivalent to 12.0±0.8 μmol NO_3_^-^-N per vial. The NO_2_^-^-N content was 0.02±0.01 mg/kg, which was equivalent to 0.04±0.02 μmol NO_2_^-^-N per vial. The NH_4_^+^-N content was 2.7±0.1 mg/kg, and the DOC content was 36.4±1.7 mg/kg. The addition dose of nitrite in the subsequent experiment was equivalent to 128.6 μmol NO_2_^-^-N per vial.

### Effect of aerobic carbon metabolism on nitrogen anaerobic reduction

Nitrate and nitrite were mainly reduced to gaseous nitrogen after 7 days of incubation in both the ANL and ANH treatments (Fig. 1a to d, Table S3). The content of NH_4_^+^-N in the UNC soil was 5.88±0.07 μmol/vial, whereas the contents of NH_4_^+^-N in the ANL and ANH treatments were only 4.62±0.45 μmol/vial and 4.17±0.19 µmol/vial, respectively, showing that no net ammonium accumulated in these two treatments. The baseline samples of the anaerobic period in the AEL and AEH treatments were AEI samples, and the content of NH_4_^+^-N was 2.01±0.50 μmol/vial in the AEI soil. However, the content of NH_4_^+^-N increased to 10.48±1.20 μmol/vial in the AEL treatment group and 27.91±0.44 μmol/vial in the AEH treatment group, indicating that the soil ammonium content significantly increased during the incubation in the AEL and AEH treatments (one-way ANOVA, P < 0.05).

**Fig. 1.**
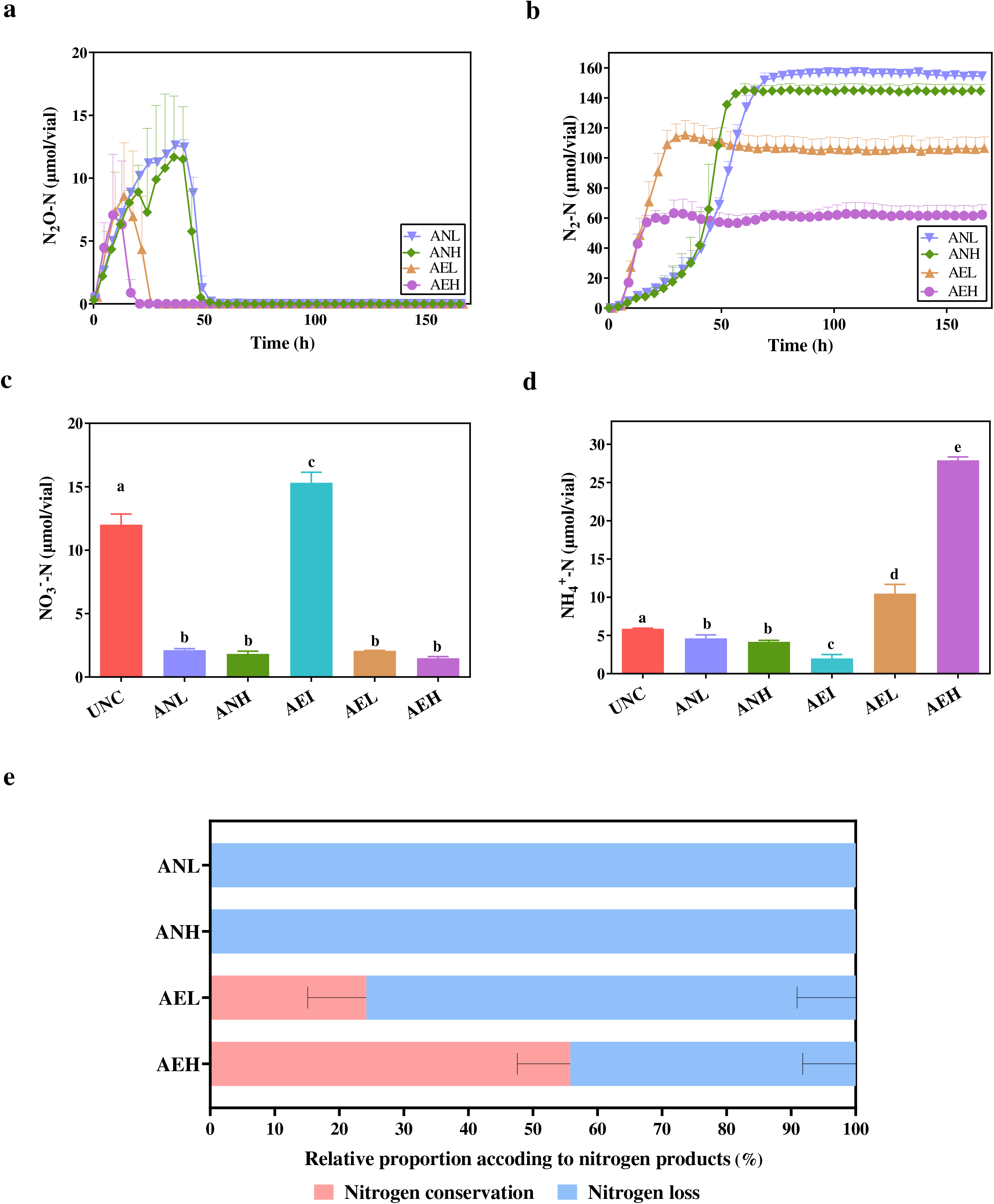
Variability in nitrogen reduction under different aeration conditions. (a) N_2_O‒N concentration dynamics. (b) N_2_‒N concentration dynamics. The data are presented as the means ± SDs in parts A and B. (c) Nitrate content of the soil samples after incubation. (d) Ammonium content of the soil samples after incubation. Different small letters indicate significant differences at the P < 0.05 level in c and d. (e) The relative proportion between nitrogen conservation and loss according to nitrogen products after incubation. All of the data are shown as the means and standard deviations in panels c to e. One-way ANOVA was used to analyze the differences among all the treatments in panels c to e.

After nitrate and nitrite were reduced by DNRA and denitrification in the different treatments, part of the nitrogen was converted into gaseous nitrogen, resulting in nitrogen loss, whereas the other part of the nitrogen was retained in the soil, resulting in nitrogen conservation. The average relative proportion of nitrogen conservation (mainly driven by DNRA in the dissimilatory reduction process) was 24.2% and 55.8% in the AEL and AEH treatments, respectively (Fig. 1e), indicating that the DNRA proportion in the AEL and AEH treatments was significantly greater than that in the ANL and ANH treatments (one-way ANOVA, P < 0.05). Conversely, nitrogen loss (mainly driven by denitrification) in the AEL and AEH treatments was significantly lower than that in the ANL and ANH treatments (one-way ANOVA, P < 0.05).

To validate the occurrence and flux of the DNRA process from nitrate to ammonium during anaerobic incubation for precluding the other ammonification, isotopic tracer incubation was performed by using ^15^N-labeled nitrate. The NH_4_^+^-^15^N atomic ratios in the AEL_14N and AEH_14N treatments (without ^15^N-labeled nitrate-amended) were 0.3687±0.0007% and 0.3687±0.0016%, respectively. Notably, the NH_4_^+^-^15^N atomic ratios in the AEL_15N and AEH_15N treatments (with ^15^N-labeled nitrate-amended) were 3.8503±1.8328% and 4.0725±1.2223%, respectively (Table 1). The NH_4_^+^-^15^N atomic ratios after the AEL and AEH treatments amended with ^15^N (AEL_15N and AEH_15N) were significantly greater (t test, P < 0.05) than those after the same treatments amended with unlabeled nitrate (AEL_14N and AEH_14N), indicating that after aerobic metabolism, a large part of the NO_3_^-^-^15^N added to the soil was reduced to NH_4_^+^-^15^N under anaerobic conditions. The results proved that the complete DNRA process from nitrate to ammonium occurred in the NCP soil after aerobic incubation with the addition of fermentable DOC.

**Table 1.** NH_4_^+^-^15^N contents in soil samples

| **Treatments** | **NH<sub>4</sub><sup>+</sup>-<sup>15</sup>N $\delta$ value** | **NH<sub>4</sub><sup>+</sup>-<sup>15</sup>N atomic ratio (atom %)** |
| --- | --- | --- |
| AEL_<sup>14</sup>N | 6.7683±2.0186 | 0.3687±0.0007 |
| AEL_<sup>15</sup>N | 9961.3127±5453.0708 | 3.8503±1.8328 |
| AEH_<sup>14</sup>N | 6.7815±4.3309 | 0.3687±0.0016 |
| AEH_<sup>15</sup>N | 10577.8556±3587.0107<sup>a</sup> | 4.0725±1.2223 |

<sup>a</sup> Data indicates mean ± standard deviation.

### Influence of incubation conditions on the nitrite ammonifiers via the *nrfA* gene analysis

High-depth sequencing analysis of the *nrfA* amplicon was performed to explore the diversity of the nitrite ammonifiers. In total, 1589,133 high-quality sequences of the *nrfA* gene were obtained from 12 samples, with an average of 132427 sequences per sample. A total of 13474 ASVs were clustered after removal of noise, chimeras and redundancy. The alpha diversity index via the *nrfA* gene showed no obvious change after incubation in any of the treatments (Table S4). There were several dominant ASVs of the *nrfA* gene (relative abundance>1%) that were abundant both in the UNC (baseline) and incubated samples, such as ASV1_Acidobacteriota, ASV2_Deltaproteobacteria, ASV3_Verrucomicrobiales, ASV4_Planctomycetota and ASV5_Deltaproteobacteria (Fig. 2a). However, some ASVs significantly responded to the specific treatments. For example, ASV9_Gemmatimonadales, ASV38_Archangium, ASV40_Bacillus, ASV97_Acidobacteriota and ASV166_Opitutaceae responded to the AEI, AEL and AEH treatments, respectively, as determined by LEfSe (Fig. 2b and Fig. S2). The identified taxonomies of these key ASVs, including Acidobacteriota, Deltaproteobacteria, Verrucomicrobiales, Planctomycetota, Gemmatimonadales, Archangium, Bacillus and Opitutaceae.

**Fig. 2.**
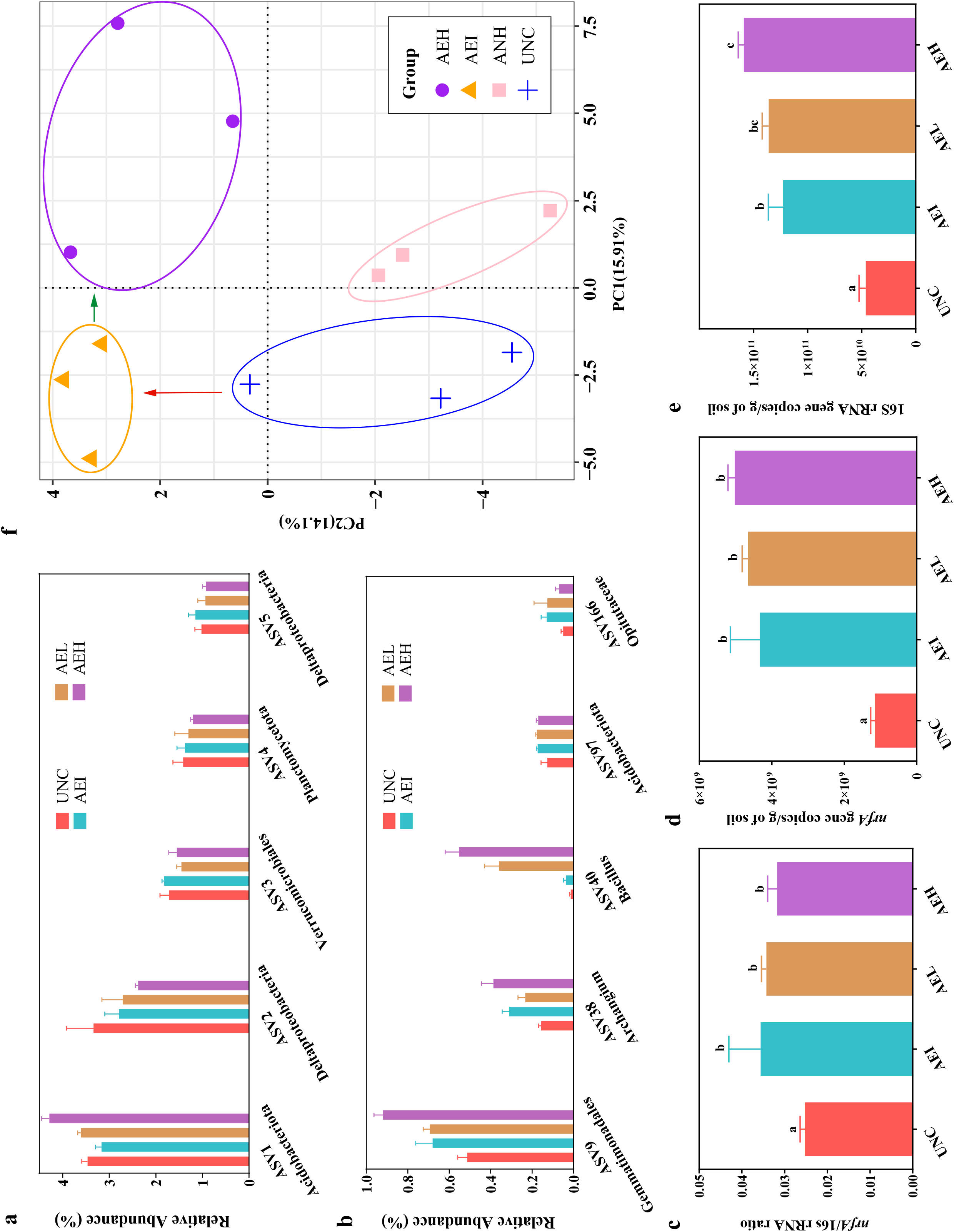
The community structure and relative abundance of nitrite ammonifiers in the samples of different treatments. (a) Relative abundance of dominant ASVs in the soil nitrite ammonifiers (relative abundance>1%) based on *nrfA* gene sequencing data. (b) Relative abundance of aerobic incubation-enriched *nrfA* ASVs based on *nrfA* gene sequencing data. (c) PCA of the nitrite ammonifiers community via *nrfA* gene species composition based on metagenomic data. PERMANOVA indicated a significant difference between the different treatments (P < 0.05). (d) *nrfA* gene copy number. (e) Ratio of *nrfA* gene copies/16S rRNA gene copies. (f) 16S rRNA gene copy number. All of the data are shown as the means and standard deviations. One-way ANOVA was used to analyze the differences among all the treatments. Different small letters indicate significant differences at the P < 0.05 level.

To understand the reason for the increase in DNRA after aerobic incubation, the abundance and composition of the *nrfA* gene were investigated in the AEI, AEL, and AEH treatments and in the uncultured control soil (UNC). The abundances of the *nrfA* gene and 16S rRNA gene were measured by quantitative PCR. The ratio of these two genes in the soil microbial community increased after aerobic incubation compared with that in the UNC samples (Fig. 2c to 2e). Bacteria containing the *nrfA* gene were significantly enriched during aerobic incubation. Afterward, in the subsequent anaerobic incubation stage, the abundances of the *nrfA* gene and bacterial 16S rRNA gene remained stable.

### Nitrogen metabolism genes reflected by metagenome sequences

In total, approximately 1.43 billion clean shotgun metagenomic reads (∼ 213.9 Gb) were generated from 12 samples, with an average of 118±9 million reads per sample. A total of 4,916,468 contigs with lengths >200 bp were assembled. In addition, 5,199,582 nonredundant ORFs were predicted and clustered. Among them, 57,562 ORFs were annotated as nitrogen cycling-related genes involved in catalyzing the processes of nitrification, denitrification, assimilatory nitrate reduction, DNRA, nitrogen fixation, and anammox. Key functional genes involved in denitrification (*narG*, *nirK*, *nirS*, *nosZ*) and DNRA (*nrfA*) were selected to reveal the functional differences between different samples. On average, 151.5±23.0 ORFs were annotated as *nrfA* genes per sample, 316.4±14.2 ORFs were annotated as *narG* genes, 829.4±51.6 ORFs were annotated as *nirK* genes, 263.5±12.2 ORFs were annotated as *nirS* genes, and 600.1±26.1 ORFs were annotated as *nosZ* genes. The other details are shown in Table S5, Fig. S3 and Fig. S4. The *nrfA* gene diversity was analyzed based on metagenomic sequencing data. The Kruskal‒Wallis test showed no significant change in the Shannon-Weaver index (P>0.05) (Fig. S5). PCA and PERMANOVA revealed that the community structure of nitrite ammonifiers changed significantly (P < 0.05) during the incubation of AEI and AEH (Fig. 2f).

The relative contributions of different genera to the total abundance of nitrogen cycling genes in the functional metagenome were calculated. The results indicated that the community compositions of the *narG* and *nrfA* genes differed among the treatments. For the *nrfA* gene, *Archangium*, *Corallococcus*, *Polyangium*, and *Sorangium* had high abundances in all the samples. However, *Bacillus* and *Sedimentibacter* were significantly enriched in the AEH treatment group. For the *narG* gene, *Nocardioides*, *Solirubrobacter*, *Microbacterium*, and *Streptomyces* had high abundances in all the samples. *Schinkia* and *Nonomuraea* were significantly enriched in the ANH treatment group. *Nocardiodes* not only accounted for a high proportion of the species composition of *narG* but also had a high relative abundance in the species composition of *nirS*, *nirK* and *nosZ* gene-carrying bacteria, indicating that *Nocardiodes* could be important for regulating the denitrification process in these soil samples (Fig. 3).

**Fig. 3.**
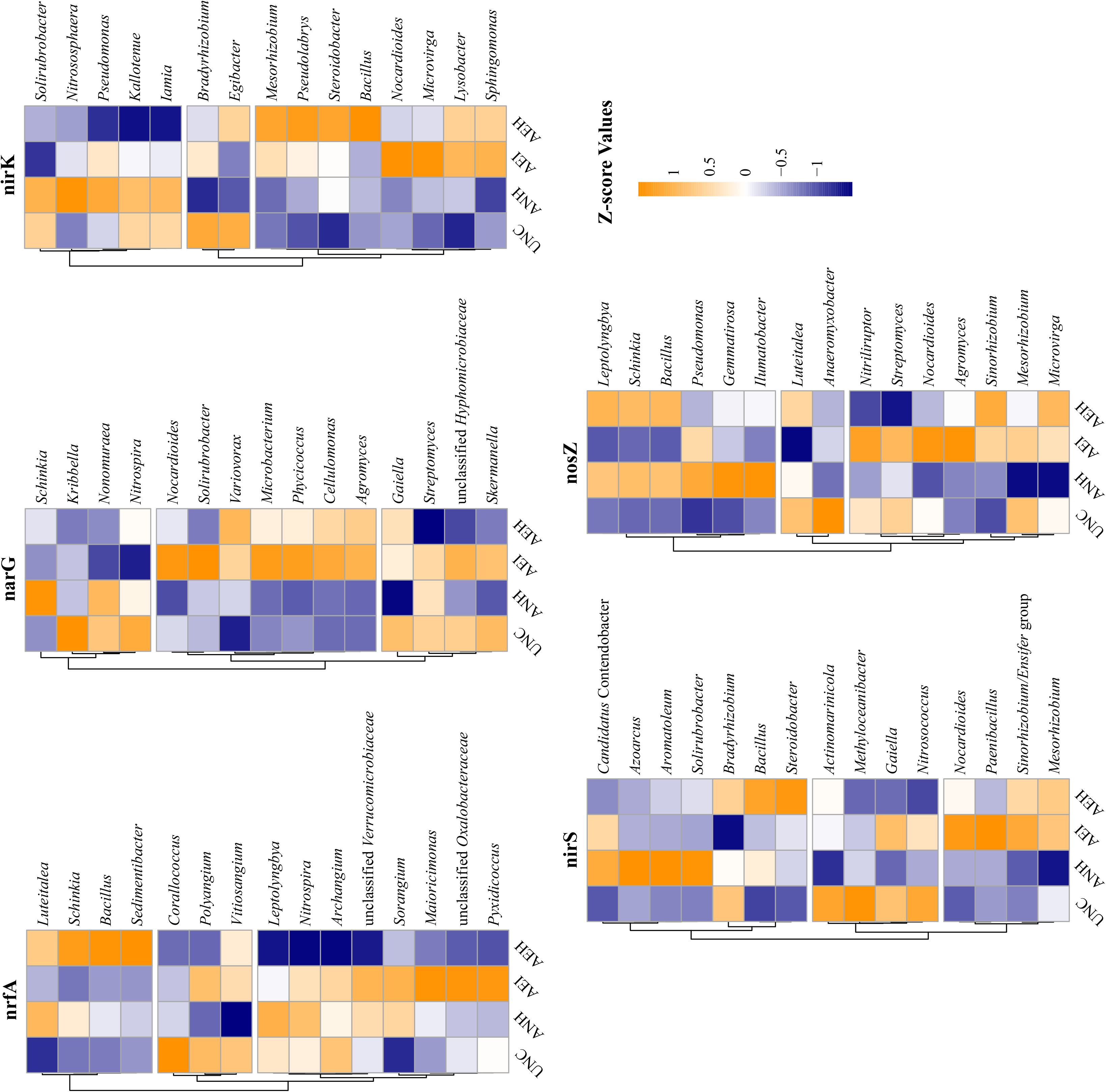
Heatmap of the standardized TPM gene aundance, showing the dominant taxonomic phylotypes of genes related to denitrification and the DNRA pathway extracted from the metagenomic data. Only the top 15 genera with different functional genes are shown here.

Phylogenetic analysis revealed that *nrfA* genes belonging to *Firmicutes* were significantly enriched in the AEH group (Fig. 4). To further explore the differences between all treatments at a finer classification level, LEfSe analysis was used to identify the key species responsible for the differences in DNRA function between the samples directly anaerobically incubated and those aerobically preincubated before anaerobic incubation. Compared with those in the UNC and ANH groups, *Bacillus marasmi*, *Serpentinicella alkaliphile* and *Planctomycetes* bacterium GWB2_41_19, which carry *nrfA* genes, were enriched in the AEI and AEH groups (Fig. S6). Additionally, seven other *nrfA*-carrying species, *Bacillus* sp. DNRA2, *Sedimentibacter* sp_B4, *Sedimentibacter saalensis*, *Myxococcaceae* bacterium, *Alkalicella caledoniensis*, *Schinkia azotoformans*, and an unclassified bacterium, were also enriched in the AEH group compared with the other groups. They might play key roles in adapting to aerobic to anaerobic conversion. The differences in the abundances of these 10 *nrfA*-carrying species among the different groups are compared in Fig. 5.

**Fig. 4.**
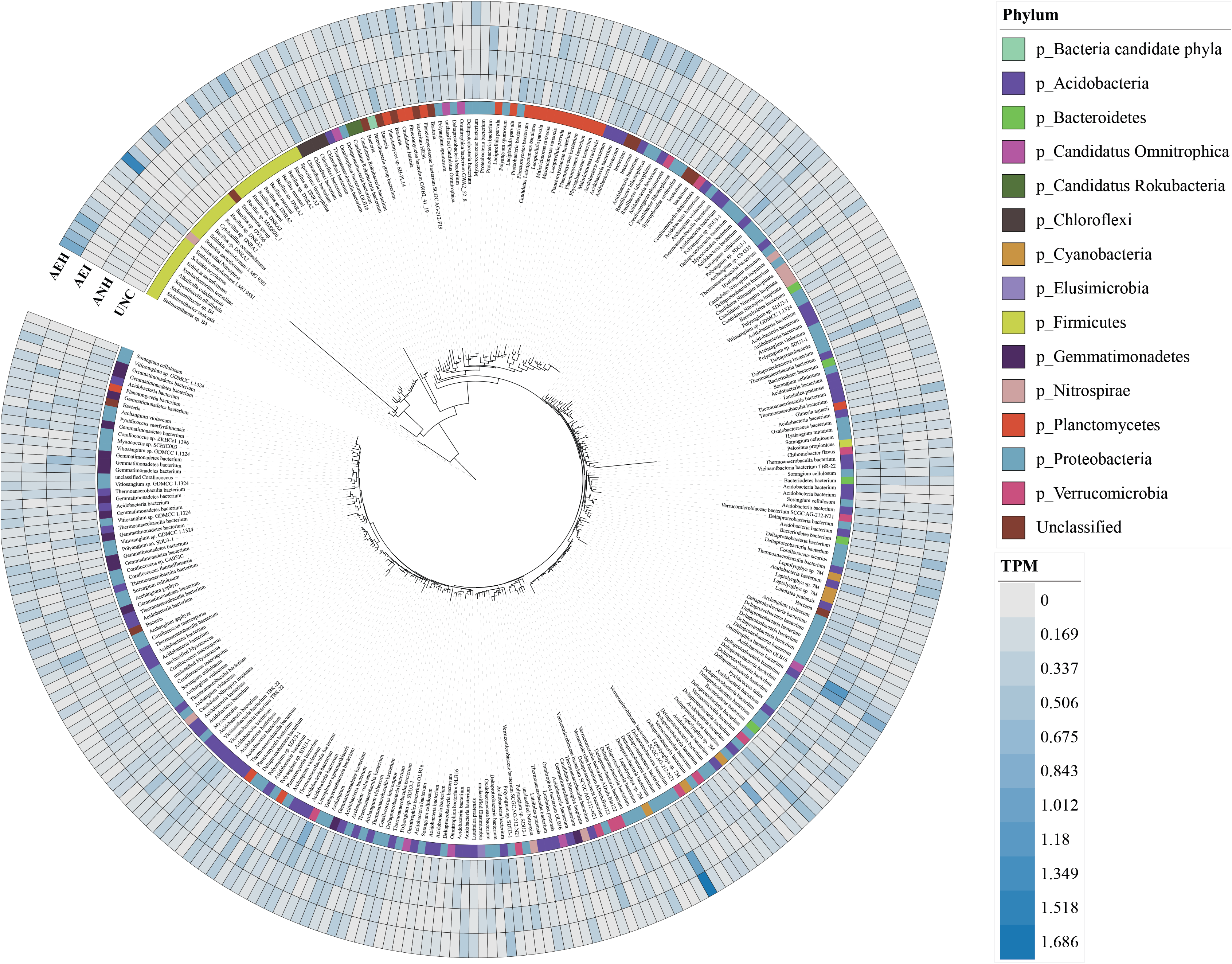
Phylogenetic inference of the *nrfA* gene according to the metagenomic data in the UNC, ANH, AEI, and AEH groups. The text in the inner circle is the highest-level species classification obtained by the annotation of different *nrfA* genes. The different colors of the middle circle attached to the phylogenetic tree represent the different phyla. The heatmap of the outer circle shows the TPM abundance of different *nrfA* genes.

**Fig. 5.**
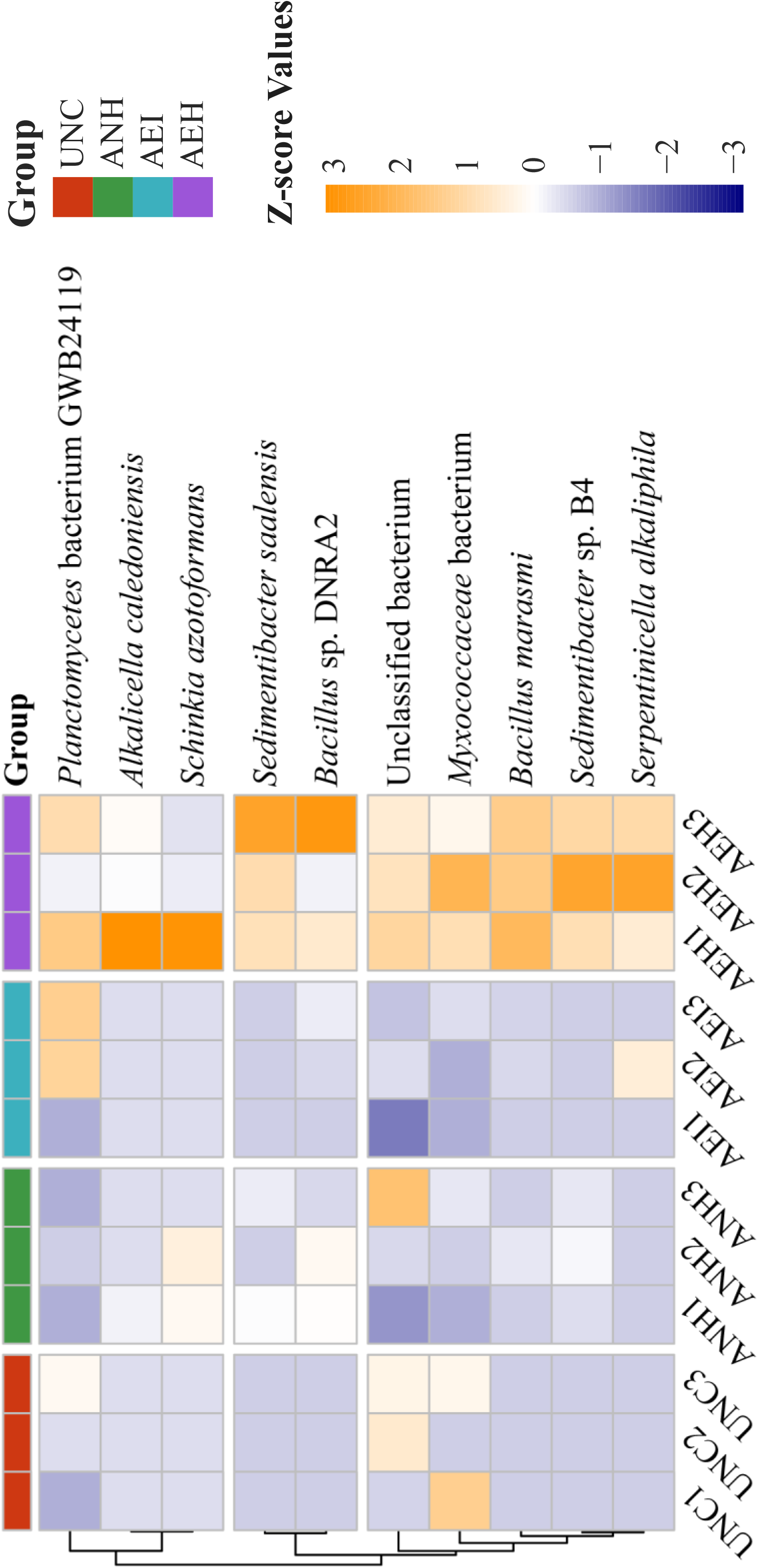
Heatmap of the standardized TPM gene aundance of 10 key nrfA-containing species (nitrite ammonifiers) enriched in the AEH treatment group according to LEfSe analysis of the metagenomic data.

## DISCUSSION

In this study, it was found that aerobic carbon metabolism modulates nitrite ammonifiers for inhibiting nitrogen loss, which was helpful for understanding soil nitrogen metabolism. Because nitrogen metabolism in the soil is very complicated due to the involvement of many metabolic processes and the various biotic and abiotic factors impacting these processes.

Moreover, there are some concerns about nitrogen metabolism in agricultural soil. First, metabolism should be beneficial or not harmful to nitrogen supplementation for plant growth. Second, nitrogen metabolism should be environmentally friendly and have a low impact on the quality of the atmosphere and underground water since nitrate is easily leached underground, and the metabolic product nitrous oxide has been recognized as a powerful greenhouse gas that has a 295-fold greater greenhouse effect than carbon dioxide. On the other hand, both denitrification and anammox processes contribute to nitrogen removal from soil (72) due to the release of gaseous products. The occurrence of DNRA could increase the ammonium content in the soil and reduce gaseous nitrogen loss due to the metabolic flux distribution between DNRA and denitrification. Therefore, the process of dissimilatory nitrate reduction to ammonium and assimilation of inorganic nitrogen to microbial biomass would be beneficial for the retention of nitrogen in the soil. Ammonium is relatively easily absorbed by plants, thus benefiting crop growth (4). Thus, the DNRA process is superior to denitrification and favors both fertilization and environmental protection in the soil ecosystem. However, enrichment of the nitrite ammonifiers in the soil is difficult due to the strong competition from the more prevalent metabolism of denitrification. In a previous study, DNRA was reported to be more pronounced in less anthropogenically influenced and nitrogen-impacted mangrove wetlands, which resulted in nitrogen conservation and a balance of nitrous oxide production (73). The predominant processes in tropical estuaries are DNRA, denitrification, and anammox (7). The function of active and competitive nitrite ammonifiers could be positively related to the carbon-to-nitrogen ratio, which has been reported in many studies (16, 26, 32, 34, 57). However, different types of agricultural soil have different nitrogen metabolism phenotypes. The qualitative and quantitative analysis of the metabolic flux of nitrogen reduction in the denitrification and DNRA pathways in this study showed the variability of nitrogen reduction in NCP soil when it was incubated under different aeration conditions.

Nitrite is the nodal intermediate metabolite in both denitrification and DNRA; therefore, we used it as a substrate to investigate the allocation between the two pathways in the microcosm experiment, intending to reduce interference from other factors. We demonstrated that the soil of a calcareous fluvo-aquic soil exclusively reduced nitrite via denitrification under constant anaerobic conditions. It has already been reported that fluvo-aquic soil possesses greater denitrification activity than other types of soils, such as black soil (31). This result raised an interesting question about how a high carbon-to-nitrogen ratio promotes DNRA and whether aeration conditions are a key factor. We performed aerobic incubations with carbon prior to anaerobic nitrite reduction to increase the abundance and composition of the nitrite reducer. Interestingly, aerobic preincubation greatly shifted nitrogen metabolism in the soil amended with labile carbon. The analysis showed that up to 55.8% of the nitrogen loss in AEH was attenuated, and nitrogen conservation was achieved when the aerobically preincubated soil was incubated under anaerobic conditions, which provides an enrichment strategy for the nitrite ammonifiers. This result indicates that the DNRA process was significantly promoted under such conditions.

To further verify the complete DNRA process in which nitrogen is reduced from nitrate to ammonium in an episode of overfertilized anoxic soil, isotope-labeled nitrate was selected as the nitrogen source substrate for stable isotopic tracer experiments. The DNRA process from nitrate to ammonium was further proven by the increase in NH_4_^+^-^15^N content. The high ratio of ^15^N measured for ammonium indicates a significantly promoted DNRA process both in AEL and in AEH, indicating that there was a strong priming effect and high organic matter (OM) mineralization during aerobic incubation, which might create appropriate conditions for DNRA facultative anaerobes (16, 25, 26, 28). Based on the nitrogen analysis and microbiome function identification, the loss of nitrate and nitrite during incubation should be mainly due to dissimilatory reduction via denitrification and DNRA processes. Here, we report that aerobic preincubation following anaerobic conditions with sufficient labile carbon could drive nitrate/nitrite reduction mainly via the process of DNRA rather than denitrification. ANH and AEH were incubated under the same carbon/nitrogen conditions, but the proportions of DNRA for nitrite reduction were distinct between the two treatments. This finding confirmed the central role of the functional bacterial community. The shifted nitrite ammonifiers community after AEI incubation in AEH and AEL showed greater competitive capacity than did the denitrifying bacterial guilds, especially under relatively high carbon conditions. According to our findings, *Sedimentibacter* was identified as one of the dominant nitrite ammonifiers in soil under anaerobic condition after aerobic carbon metabolism. Additionally, it was known that *Sedimentibacter* was also the typical bacteria in sediments as previously described (74). Besides sediments as the environments of oxic-anoxic interfaces possessed a high occurrence rate of DNRA according to previous study (38). Accordingly, it was inferred that alternative aerobic and anaerobic condition might be the suitable ecological niche for nitrite ammonifiers.

To elucidate the underlying biological and ecological mechanisms, we measured the quantity of related key functional genes and the composition of *nrfA*, a key DNRA gene. In this study, the significant increase in nitrite ammonifiers in the quantification measurement proved that the prerequisite abundance of functional microorganisms is essential for the occurrence of DNRA, as indicated in previous studies (32, 57). We also found that the functional community for the nitrite ammonifiers shifted after aerobic carbon metabolism for the soil provided with carbon. The extra preincubation significantly enriched a group of nitrite ammonifiers, as shown by *nrfA* gene sequencing and metagenome annotation. In previous studies, soil redox potential was reported to influence the frost soil microbiome and N cycling rate (75, 76). One possible reason for this phenomenon is that DNRA was less sensitive to oxygen exposure than expected, and the organisms mediating this reductive process were also tolerant of oxic conditions (76) which is consistent with our findings. In this study, the alteration of nitrite ammonifiers community structures in the treated soil could be the reason driving the occurrence of the DNRA process. In particular, the abundant nitrite ammonifiers such as *Sedimentibacter* might be the main contributors to nitrogen reduction to ammonium.

## SUPPLEMENTAL MATERIAL

SUPPLEMENTAL MATERIAL.docx, 1074 KB.

Supplementary Results, Table S1 to S5, Figure S1 to S6

## DATA AVAILABILITY

The *nrfA* gene fragment sequences were submitted to the Sequence Read Archive (SRA) database of the National Center for Biotechnology Information (NCBI) under the accession number SRP291604. The metagenomic sequences were submitted to the Sequence Read Archive (SRA) database of the National Center for Biotechnology Information (NCBI) under the accession number SRP395973.

## ACKNOWLEDGMENTS

This work was supported by the National Natural Science Foundation of China (NSFC 31971526) and the National Key Research and Development Program of China (2017YFD0200102). The authors would like to express their gratitude to Prof. Xiaotang Ju for his help in collecting the soil samples.

## AUTHOR CONTRIBUTIONS

Xiaogang Wu: Investigation, Software, Methodology, Data curation, Formal analysis, Visualization, Writing – original draft, Writing – review & editing. Siyu Yu: Investigation, Software, Methodology, Data curation, Visualization, Writing – original draft, Writing – review & editing. Weikang Sui: Investigation, Methodology, Data curation, Validation, Writing – original draft. Xinyu Zhang: Methodology, Data curation. Ji Li: Methodology, Data curation. Qiaoyu Wu: Methodology. Xiaojun Zhang: Resources, Conceptualization, Writing –review & editing, Supervision, Project administration, Funding acquisition.

## CONFLICTS OF INTEREST

The authors declare that they have no competing interests.

